# Benchmark data and software for assessing genome-wide CRISPR-Cas9 screening pipelines

**DOI:** 10.1101/2022.09.23.509258

**Authors:** Raffaele Iannuzzi, Ichcha Manipur, Clare Pacini, Fiona M. Behan, Mario R. Guarracino, Mathew J. Garnett, Aurora Savino, Francesco Iorio

## Abstract

Genome-wide recessive genetic screens using lentiviral CRISPR-guide RNA libraries are widely performed in mammalian cells to functionally characterise individual genes and for the discovery of new anti-cancer therapeutic targets. As the effectiveness of such powerful and precise tools for cancer pharmacogenomic is emerging, reference datasets for their quality assessment and the validation of the underlying experimental pipelines are becoming increasingly necessary. Here, we provide a dataset, an R package, and metrics for the assessment of novel experimental pipelines upon the execution of a single calibration viability screen of the HT-29 human colon cancer cell line, employing a commercially available genome-wide library of single guide RNAs: the Human Improved Genome-wide Knockout CRISPR (Sanger) Library. This dataset contains results from screening the HT-29 in multiple batches with the Sanger library, and outcomes from several levels of quality control tests on the resulting data. Data and accompanying R package can be used as a toolkit for benchmarking newly established experimental pipelines for CRISPR-Cas9 recessive screens, via the generation of a final quality-control report.

## Background & Summary

Genome-wide CRISPR-Cas9 screens are being increasingly employed to explore various genotype–phenotype associations^1^, to identify genes whose function is essential for cell viability and proliferation (essential genes or fitness genes), and new potential targets for personalised anti-cancer therapies^2–7^. Several methods exist for assessing the quality of the datasets derived from these screens, evaluating sequence quality, single-guide RNA (sgRNA) count distributions and negatively selected genes^8^. In addition, comprehensive analyses have been performed to evaluate the level of reproducibility and integrability of large-scale cancer dependency datasets assembled from independently performed CRISPR-Cas9 screens^9,10^. However, to date no easy-to-use toolkit is available to assist experimental scientists in validating newly established experimental pipelines for genome-wide CRISPR-Cas9 genetic screens using pooled sgRNA libraries.

In Behan *et al*.^7^, we performed genome-wide CRISPR–Cas9 fitness screens of 339 cancer cell lines from the Cell Models Passport panel^11^. We analysed the data resulting from this screen with an ad-hoc computational pipeline designed to identify new anti-cancer therapeutic targets at a genome-scale. To this aim, we defined quality control assessment practices and applied stringent quality control criteria, finally retaining data for 324 cell lines. Via a target-prioritisation bioinformatics pipeline we predicted and validated a novel selective therapeutic target for cancers with microsatellite instability: the Werner syndrome ATP-dependent helicase^7^ (a finding simultaneously reported by other independent studies^12–14^). Results and datasets from this study are available for download on the Project Score data portal (https://score.depmap.sanger.ac.uk/). As part of this effort, we screened the HT-29 colorectal cancer cell line with the same experimental settings in multiple batches and dates, to assess robustness and reproducibility of our experimental pipeline.

Here, we provide high-quality data from 30 screens of the HT-29 cell line yielding reliable gene essentiality profiles, and a dedicated analytical tool implemented into an R package^15^. We propose the use of this data and software as a simple toolkit to benchmark and validate newly established genome-scale CRISPR-Cas9 knock-out screening pipelines employing the Human Improved Genome-wide Knockout CRISPR sgRNA library (the Sanger library, available on Addgene)^16^. By performing a single calibration screen of the HT-29 cell line with the Sanger library and settings described in Behan *et al*.^7^, experimental scientists can assess quality and reproducibility of their pipeline by processing resulting data with our R package, which implements a diversified set of metrics to compare new data with expected outcomes.

Data and code, including the *HT29benchmark* R package, are available at https://score.depmap.sanger.ac.uk/ downloads, FigShare^17^ and https://iorio-apps.fht.org/.

## Methods

### Reference dataset generation: CRISPR-Cas9 screens

The protocol used for the generation of Cas9-expressing HT-29 cell lines and transduction of the Sanger library^16^ is described in Behan *et al*.^7^. Briefly, we employed the commercially available Sanger Library v1.0 (Addgene, 67989), encompassing 90,709 sgRNAs targeting 18,009 genes, and a second version of the same library (Sanger library v1.1) including all the sgRNAs from v1.0 plus 1,004 non-targeting sgRNAs, and 5 additional sgRNAs targeting 1,876 selected genes encoding kinases, epigenetics-related proteins and pre-defined fitness genes, for a total of 10,381 additional sgRNAs. Cells were grown for 14 days following transduction with the Sanger Library (v1.0 or v1.1) and selection, and then collected for DNA extraction. Illumina sequencing and sgRNA counting were performed as described in Tzelepis *et al*^16^. Experiment identifiers and settings are fully described in Supplementary Table 1, summarised in Table 1 and further detailed in the Extended data Fig. 2j of Behan *et al*.^7^.

**Table 1.** Reference HT-29 screening dataset. Libraries, experiment identifiers and transfection/selection efficiencies across screens.

| Sanger library version | Experiments identifiers | N. of replicates | Cas9 activity | Average transfection efficiency | Average Puromycin selection |
| --- | --- | --- | --- | --- | --- |
| 1.0 | HT29_c903 | 6 | 94.8% | 2.33% | 83.53% |
| 1.0 | HT29_c904 | 3 | 94.8% | 27.57% | 89.97% |
| 1.1 | HT29_c905 | 9 | 94.8% | 33.42% | 80.81% |
| 1.1 | HT29_c906 | 6 | 94.8% | 35.65% | 88.40% |
| 1.1 | HT29_c907 | 3 | 94.8% | 32.40% | 89.07% |
| 1.1 | HT29_c908 | 3 | 94.8% | 32% | 79.33% |

Overall, we performed 2 independent experiments with the Sanger v1.0 library and 4 experiments with the Sanger v1.1 library. These can be regarded as biological replicates of HT-29 CRISPR screens, while each experiment has been performed with a varying number of technical replicates (from 3 to 9) for a total of 30 individual screens, as indicated in Table 1.

### Reference dataset preprocessing

We quantified and pre-processed post library-transduction and control library-plasmid sgRNA read counts as described in Behan *et al*.^7^, removing sgRNAs with less than 30 reads in the library-plasmid and keeping only sgRNAs in common between the two versions of the Sanger libraries. Subsequently, we normalised counts across replicates, scaling each sample by total number of reads. Post normalisation, we computed sgRNA log fold-changes () between individual replicate read counts and library-plasmid read counts for each experiment, keeping the replicates separated (Supplementary Fig. 1). These pre-processing steps were performed with the `ccr.NormfoldChanges` function of our previously published *CRISPRcleanR* R package^18^, using default parameters. Resulting data at all intermediate pre-processing levels are included in our reference dataset (available at: https://score.depmap.sanger.ac.uk/downloads and on FigShare^17^).

### Example of user provided data

In order to demonstrate and test the diverse functionalities of the *HT29benchmark*R package, we used (as an example of user-provided data) a lower quality screen of the HT-29 cell line, which was discarded from the analysis set in Behan *et al*.^7^, and encompasses six technical replicates of an HT-29 screen, obtained following the same screening protocol and the pre-processing steps described above.

### Receiver operating characteristic analysis

To compute receiver operating characteristic (ROC) and precision/recall (PrRc) curves, required to perform high-level quality control assessment of CRISPR-cas9 screens, we used the `HT29R.individualROC` function of the *HT29benchmark*R package, which implements the `ROC_Curve` and `PrRc_Curve` functions of the *CRISPRcleanR* package^18^ (version 2.2.1), which itself implements the `roc` and `coords` functions of the *pROC* open-source R package (version 1.18.0)^19^.

### Fitness-effect threshold

Following the approach we presented in Pacini *et al*.^10^, we employed a rank-based method to compute a fitness effect significance threshold for each HT-29 reference screen, thus identifying a set of significantly depleted (or essential) genes at a fixed level of 5% false discovery rate (FDR), based on their depletion log fold-changes (LFCs). Specifically, in a given screen, we first ranked all genes in increasing order of average depletion LFCs (based on the differential abundance of their targeting sgRNAs at the end of the assay versus plasmid control). Then we scrolled the obtained ranked list from the most depleted gene to the least depleted one, and we considered the depletion LFC *r* of each encountered gene as a potential threshold, i.e. calling all genes with a depletion log fold-change *< r* significantly depleted.

Among the significantly depleted genes at a candidate threshold *r* we focused only on those belonging to any of two prior known sets of essential (*E*) and non-essential (*N*) genes^2^. Considering these two sets as reference positive and negative controls, respectively, allowed us to compute a positive predictive value (PPV), thus a false discovery rate (FDR = 1 − PPV). We finally select as fitness-effect significance threshold the largest *r*, yielding an FDR 0.05. We implemented this procedure using the `roc` and `coords` functions of the *pROC* open-source R package (version 1.18.0)^19^ implemented in the `HT29R.ROCanalysis` and `HT29R.FDRconsensus` functions of the *HT29benchmark* R package.

### Data visualisation

For data visualisation, we used R base graphics plus the following R libraries and packages (listed in alphabetical order), all available on bioconductor^20^ or on The Comprehensive R Archive Network (CRAN) repository: *crayon* version 1.5.1; *enrichPlot* version 1.14.2; *GGally* version 2.1.2; *ggplot2* version 3.3.6; *ggrastr* version 1.0.1; *grid* version 4.1.0; *gridExtra* version 2.3; *gtable* version 0.3.0; *RcolorBrewer* version 1.1.3; *VennDiagram* version 1.7.3; *vioplot* version 0.3.7^21^.

### Enrichment analysis

We performed Gene Ontology (GO) enrichment analysis to identify biological processes over-represented in the list of HT-29-specific fitness genes. For this analysis, we used the *org*.*Hs*.*eg*.*db* R package (version 3.14.0) to retrieve the gene universe and the *clusterProfiler* R package (version 4.2.2) to perform the enrichment analysis of the HT-29-specific genes.

### Data Records

The entire HT-29 reference dataset described here is available at different intermediate levels of pre-processing on the Project Score website https://score.depmap.sanger.ac.uk/downloads and on FigShare^17^ (https://figshare.com/articles/dataset/HT29_reference_dataset/20480544).

The main data folder contains four subfolders:

- **00_rawCounts assembled** - Containing one tsv file for each HT-29 screen. Each file comprises the control library-plasmid sgRNA counts, as well as 14 days post-selection sgRNA counts across technical replicates;
- **01_normalised_and_FCs** - Containing Rdata files of normalised counts and depletion Log fold-changes (LFCs) for the six screens, plots of counts’ distribution pre- and post-normalisation, and boxplots showing LFCs’ distributions (PDF files);
- **02_lowLev_QC** - subdivided in the following four subfolders:
  1. FC_distr - log fold-change distribution plots for each of the six screens, in PDF;
  2. FC_Rep_corr - Between-replicates correlation plots for each of the six screens, in PDF;
  3. PrRc_curves_ind_rep - Plots of replicate Precision Recall (PrRc) curves quantifying essential/non-essential genes’ classification performances across the six screens, in PDF;
  4. ROC_curves_ind_rep - Plots of replicate Receiver Operating Characteristic (ROC) curves quantifying essential/non-essential genes’ classification performances across the six screens, in PDF;
- **03_HL_QC_Stats** - Density plots of depletion LFCs for reference gene sets across the six experiments with quality control values, in PDF.

### Technical validation

In the *HT29benchmark* package we have implemented a set of reference metrics for the assessment of quality and reproducibility of CRISPR screens. In particular, these metrics assess sgRNA LFC distributions, screen outcomes’ reproducibility across technical replicates, inter-screen similarity, and screens’ ability to detect known fitness genes among the depleted ones. Here, we report results from applying these metrics to technically validate our HT-29 reference dataset, as well as to showcase how our package can be used to evaluate an example of user-provided dataset. Furthermore, we report a set of reliable HT-29 specific fitness genes, which we have identified via a joint analysis of all the screens in our reference dataset. These genes are expected to be detected as significantly essential in any CRISPR screen of the HT-29 cell line performed with the experimental settings underlying the generation of our reference dataset^7^, and using the Sanger library^16^. All the technical validations presented here can be re-executed by a user on its own data through our *HT29benchmark*R package.

#### *HT29benchmark*R package overview

The *HT29benchmark*R package allows assessing quality and reproducibility of both reference and user-provided CRISPR screens of the HT-29 cell lines employing the Sanger library and the experimental settings described in Behan et al^7^. More in detail, the *HT29benchmark*package implements several routines from our previously published *CRISPRcleanR* package^18^ wrapped in novel ad-hoc designed functions, providing a powerful and easy-to-use tool able to:

- Download the HT-29 reference dataset;
- Inspect and visualise sgRNAs depletion LFC distributions of each screen;
- Evaluate intra-screen reproducibility of depletion LFCs at the sgRNA level, as well as at the gene level;
- Evaluate inter-screen similarity of depletion LFCs at the sgRNA level, as well as at the gene-level;
- Evaluate individual screen performances in correctly partitioning known essential (positive control) and known non-essential (negative control) genes, when considered as rank based classifiers based on gene depletion LFCs - through ROC and PrRc curves, as well as Recall at a fixed False Discovery Rate (FDR);
- Visualise depletion LFC distributions for positive and negative control genes (as well as for their targeting sgRNAs) and compute Glass’s D scores quantifying the difference of their average depletion LFCs in the screen under consideration;
- Derive HT-29-specific essential/non-essential genes, by analysing all screens in the reference dataset jointly and then to use these sets as positive/negative controls to estimate to what extent a user-provided screen meets expectations, based on the metrics listed above.

### Inspection of sgRNA log fold-change distributions

The `HT29R.FCdistributions` function of the *HT29benchmark* package allows inspecting sgRNA LFC distributions and it computes statistics such as average range, median, interquartile range, 10^th^ − 90^th^ percentile range, skewness and kurtosis. We have applied these metrics to the screens in our reference HT-29 dataset, observing that the LFC distributions and their parameters meet expected shape/values of a typical CRISPR-Cas9 recessive screen^22,23^ (Fig. 1a).

**Figure 1.**
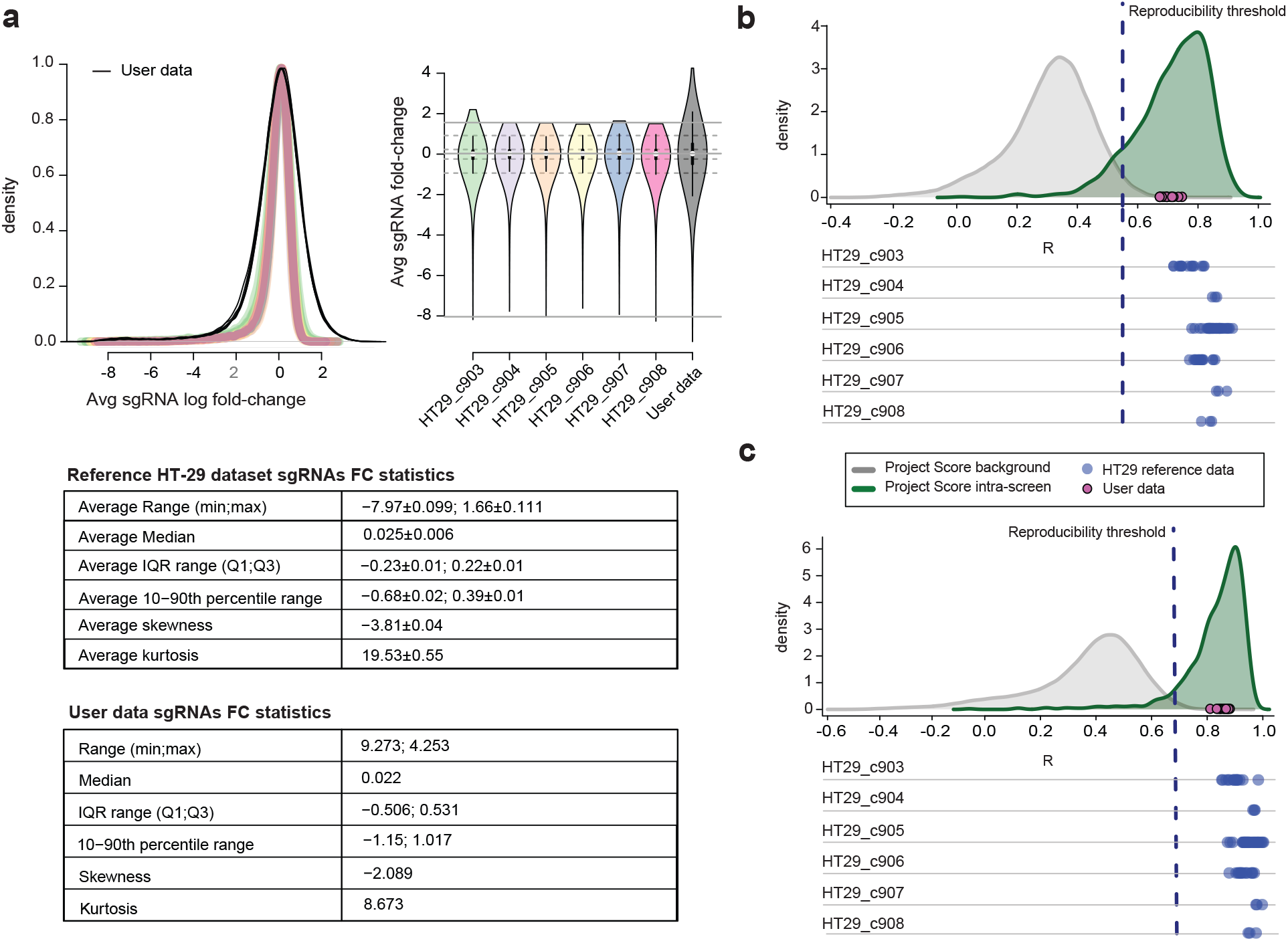
(a) Distributions of single-guide RNA (sgRNA) depletion log fold-changes and their average parameters (with confidence intervals) across the different screens of the reference HT-29 dataset, and in an example of user-provided screen performed using reagent and experimental settings described in Behan *et al*.^7^ and the Sanger library. (bc) Outcomes from an evaluation of inter-screen similarity. Distributions of pairwise Pearson’s correlation scores computed between gene essentiality profiles of replicates for each of the six HT-29 reference screens (blue dots), considering depletion log fold-changes of highly reproducible/informative sgRNAs only. Their value is abundantly larger than the quality control threshold defined from the analysis of the Project Score dataset (dark blue dashed vertical line), both at sgRNA- (b) and gene-level (c). The distribution of correlations from comparing replicates of the same screen in Project Score is shown in green, while the distribution of correlations from comparing each possible pair of replicates (regardless the screen) is shown in grey, with densities varying according to the level inspected (sgRNA or gene). The magenta points indicate correlation between pairs of replicates of an example user-provided screen of the HT-29 cell line (performed using the same setting of Behan *et al*.^7^) and the Sanger library^16^ which in this case exceeds the reproducibility threshold.^16^.

This function can also take in input a user-provided screen, allowing a comparison between reference and new data, which might unveil unexpected distribution shapes, outliers and other data inconsistencies, thus allowing a first exploratory assessment of a new screen (Fig. 1a).

### Intra-screen reproducibility assessment

To assess screen replicates’ reproducibility, we defined a reliable measure of intra-screen similarity. In our previous work^7^, we observed that comparing replicates of the same screen at the level of absolute post-transduction sgRNA count profiles produces meaningless outcomes, due to individual sgRNA counts varying in different ranges, which are determined by their initial amount in the library-plasmid. This produces a strong Yule-Simpson effect^24^ resulting in a generally high background correlation between any pair of genome-wide sgRNA count profiles. As a result, when using this criterion as a reproducibility metric, pairs of replicates of the same screen are indistinguishable from two individual replicates of different screens (Supplementary Fig. 2a).

Due to only a small fraction of genes having an impact on cellular fitness upon CRISPR-Cas9 targeting, pairs of replicates from different screens tend to yield generally highly correlated dependency profiles even when considering sgRNA (or gene level) depletion LFCs (Supplementary Fig. 2bc) instead of absolute counts.

For these reasons, in Behan *et al*.^7^ we followed an approach similar to that introduced in Ballouz *et al*.^25^ and identified a set of library-specific informative, and highly reproducible, sgRNAs pairs targeting the same gene and with an average pairwise correlation of their depletion LFC pattern greater than 0.6 across a set of 332 cell lines from Project Score^7^. This yielded a total of 838 unique informative sgRNAs. Per construction, the depletion patterns of these sgRNAs are both reproducible and informative, as they involve genes carrying an actual and sufficiently variable fitness signal.

When considering these informative sgRNAs only, correlation scores from comparing replicates of the same screens were significantly higher than those from comparing pairs of replicates from different screens (Supplementary Fig. 2de) of the Project Score dataset. This allowed us to define a threshold value discriminating the two distributions both at sgRNA- and gene-level (R = 0.55 and R = 0.68, respectively), as defined in Behan *et al*.^7^ (Fig. 1bc), and to use this value as a required minimal quality while evaluating intra-screen reproducibility.

The function `HT29R.evaluateReps` of the *HT29benchmark* package allows a robust assessment of input screens, producing plots like those shown in Fig. 1bc. All technical replicate pairs in the HT-29 reference screens exceed the reproducibility threshold defined in Behan *et al*.^7^ (blue circles in Fig. 1bc). Moreover, inter-screen reproducibility of user-provided data can also be evaluated (magenta circles in Fig. 1bc), and results visualised and compared with those observed for the reference HT-29 dataset.

### Inter-screen similarity evaluation

As a second measure of reproducibility, we evaluated results’ comparability across different screens. Thus, we considered genes (or sgRNAs) passing pre-processing filters in all the six HT-29 screens, computed LFCs’ profiles and averaged them across technical replicates, ending up with six different LFC profiles (one for each screen). We computed Pearson’s correlation scores comparing each pair of these profiles. This analysis is performed (and results can be visualised) by the `HT29R.expSimilarity` function included in our *HT29benchmark* package, which (as before) can be also used on a user-provided screen to assess its similarity, in terms of depletion LFCs, to the six HT-29 reference screens. For consistency with the reproducibility measure introduced in the previous section, this function allows considering the entire Sanger library or highly informative sgRNAs only, and to evaluate screens’ similarity both at the sgRNA and gene level (Fig. 2 and Supplementary Fig. 3abc).

**Figure 2.**
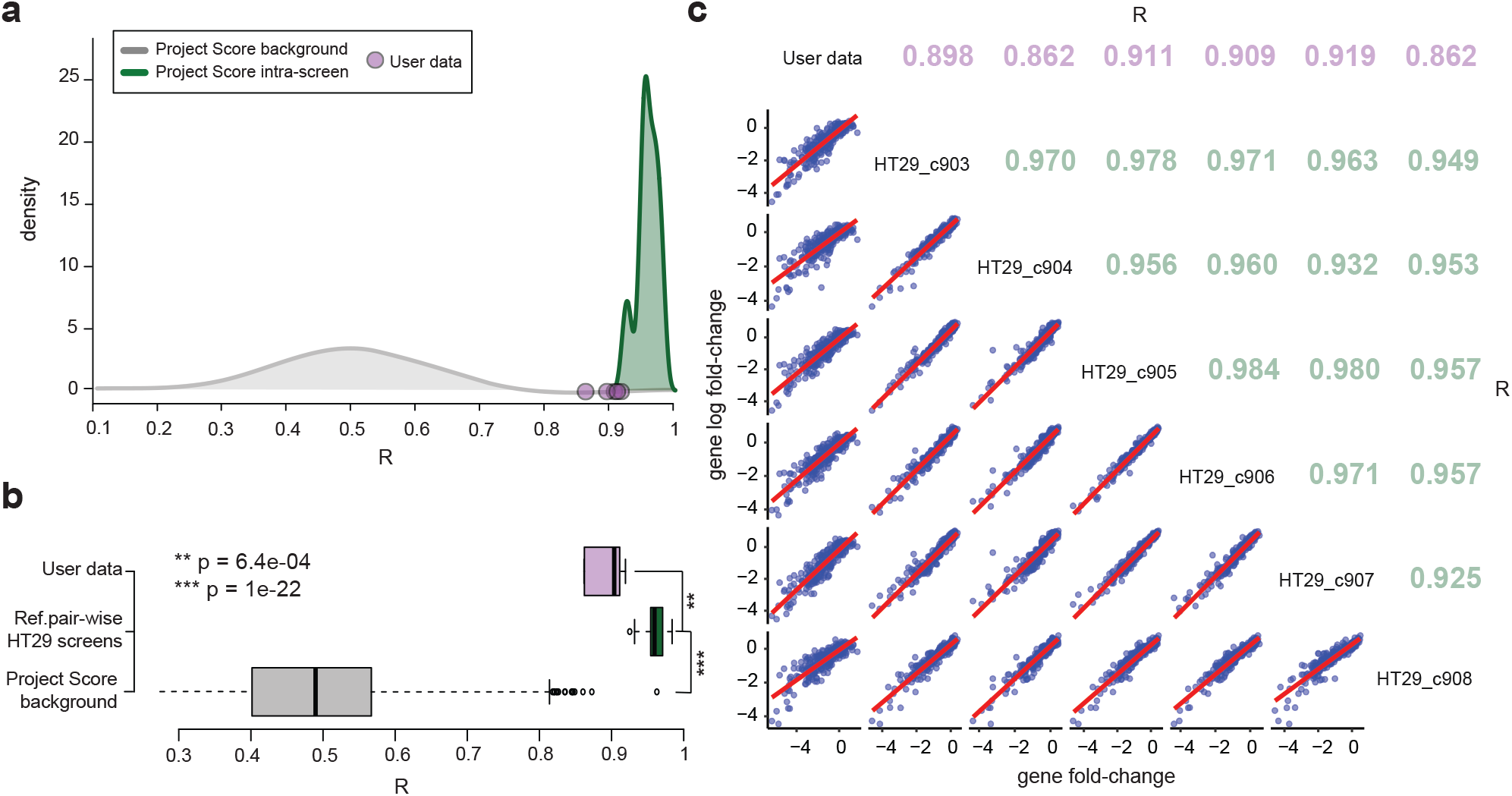
Inter-screen similarity evaluation. (a) Pearson’s correlation scores between profiles of depletion log fold-changes (LFCs) computed at the gene level using the subset of reproducible and highly informative sgRNAs (n = 838) between pairs of HT-29 screens (in green) and between the HT-29 reference screens and an example user-provided screen (in pink), with replicates collapsed by LFC averaging. The distribution in grey are computed between each possible pair of screen replicates in Project Score, to estimate expectation is also visualised (in gray). (b) Two-sided t-test comparing expected Project Score correlation scores versus those computed between each pair of screens in the HT-29 reference dataset, as well as those computed between the example data screens versus those computed in the HT-29 reference dataset. The reference dataset scores are largely significantly different from expectation, the user data scores are still largely different from expectation but not as much as the reference data. (c) Scatter-plot correlation matrix showing pairwise Pearson’s correlation scores computed within HT-29 references and between user data versus HT-29 reference screens.

### Screens classification performances

The ability to discriminate prior known essential and non-essential genes based on their depletion LFC observed in a CRISPR-Cas9 recessive screen is widely used to assess the quality of that screen^3,5,7,9,10,23,26,27^.

In particular, a good quality CRISPR screen will tend to detect genes involved in fundamental cellular processes, and other *core fitness genes*, as highly depleted invariantly across screened cell types. Robust reference sets of core essential and non-essential genes can be used as a gold standard to evaluate screens’ performances^26,28^. The `HT29R.PhenoIntensity` function provides a measure of screen quality by leveraging the intensity of the phenotype exerted by inactivating these genes. To quantify this effect, in Behan *et al*.^7^ we computed Glass’s D score^29^ computed respectively for reference essential genes (i.e., genes that reduce cellular viability/fitness upon inactivation^3^) (*E*) and (more stringently) for ribosomal protein genes^30^ (*R*) genes. These scores account for the difference between the average depletion LFCs of the genes in *E* (respectively *R*) and that of genes known to be non-essential^3^ (*N*) in relation to the standard deviation of the depletion LFCs of the genes in *E* (respectively *R*), as it follows:

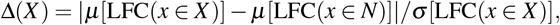

where *X ∈ E, R*, and *μ* and *s* indicate mean and standard deviation, respectively. The Δs for the screens in the reference dataset were consistently *>* 2 for ribosomal protein genes and *>* 1 for the other essential genes (with a Glass’s *Delta >* 0.8 widely considered an indicator of large effect size), thus indicative of generally good data quality (Fig. 3a and Supplementary Fig. 4). In addition, as depicted in Fig. 3ab, in this case applying this metric to the example user-provided screen yielded values within the expected ranges.

**Figure 3.**
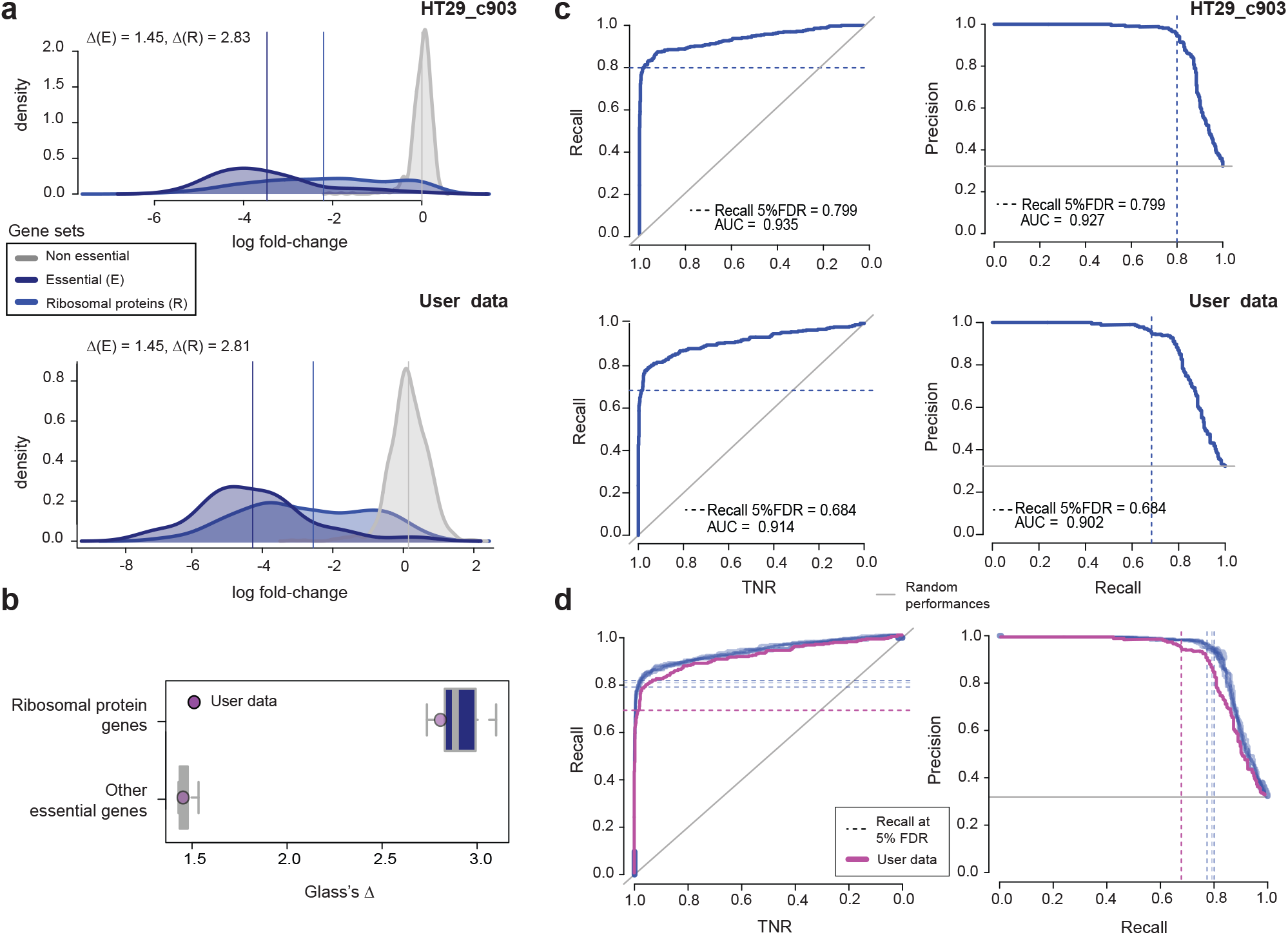
Screens’ quality in terms of phenotype Intensity and receiver operating characteristic (ROC) analysis. (a) Distributions of gene depletion log fold-changes (LFCs) for one of the screens in the HT-29 reference dataset (at the top) and an example user-provided screen (at the bottom). Glass’s D (GD) scores for reference essential genes (*E*) and ribosomal protein genes (*R*) with respect to non-essential genes are reported at the top of each plot. Vertical lines indicate mean LFCs for each gene set as indicated by the different colours. (b) Distributions of GD scores with respect to ribosomal protein genes and other essential genes (as indicated by the different colours), computed across the reference screens with overlaid GΔs observed for the example user-provided screen. (c) ROC and Precision Recall (PrRc) curves quantifying the ability of a given screen in correctly classifying prior known essential and non-essential genes, based on their depletion LFCs for one of the screens in the HT-29 reference dataset (at the top) and an example user-provided screen (at the bottom). Recall of prior known essential genes at a 5% false discovery rate and areas under the curves are also reported, with the former indicated also by the dashed lines. (d) As for panel c but extended to all the reference screens and the user data, as indicated by the different colours.

In addition to the Glass’s Δs, we implemented and included in our package the `HT29R.ROCanalysis` function computing and visualising ROC and PrRc curves to evaluate the ability of each screen in correctly partitioning prior known essential (*E*) and non-essential (*N*) genes, when considered as a rank based classifier based on sgRNA- or gene-depletion-LFCs (as explained in the previous sections). Applying this function to the HT-29 reference dataset, as well as to example user-provided data, yielded the results shown in Fig. 3cd. Also in this case our reference dataset yielded very good quality scores.

Finally, as a further quality assessment and reference to the user, we computed fitness effect significance thresholds employing prior known essential and non-essential genes^31^ at different FDR levels, and we quantified corresponding Recall values of prior known essential-genes, as well as a novel set of human core-fitness genes introduced in Behan et al^7^ and various sets of other essential genes (all available in the *CRISPRcleanR* package^18^(Supplementary Fig. 5 and Supplementary Table 2). Also these results confirmed the high quality of our reference dataset.

### HT-29-specific fitness genes

We assembled a list of genes that are consensually significantly depleted across all our reference HT-29 screens, thus should be observed as significantly depleted in new screens of the HT-29 cell line performed with the Sanger library^16^ and the experimental setting described in Behan *et al*.^7^. First of all, for each reference HT-29 screen we identified a set of genes significantly depleted at a 5% FDR and its complement, i.e. a set of genes not significantly depleted, using reference sets of essential (*E*) and non-essential (*N*) genes^31^ to compute significance thresholds, as explained in the previous sections. Intersecting all these sets of screen-specific significantly depleted, respectively non depleted, genes yielded a high-confidence set of HT-29 specific essential, respectively non-essential, genes. We assessed how each reference screen discriminated these two sets in terms of Glass’s Δ^29^ or Cohen’s d^32^, computed as explained in the previous sections. This allowed us to define again a set of expected values to evaluate a newly performed screen of the HT-29 (Fig 4abc). These HT-29-specific fitness genes are also provided in Supplementary Table 3, partitioned into three tiers based on their average depletion LFCs across screens. These genes showed a fairly consistent depletion LFCs across screens (Fig 4d) and were significantly enriched for previously report human essential genes (Fisher’s exact test *p* = 7.1 * 10 − 221, Fig. 4c) and for fundamental biological processes (BP) such as “ribosome biogenesis” and “RNA splicing” (Fig. 4e), confirming their reliability.

**Figure 4.**
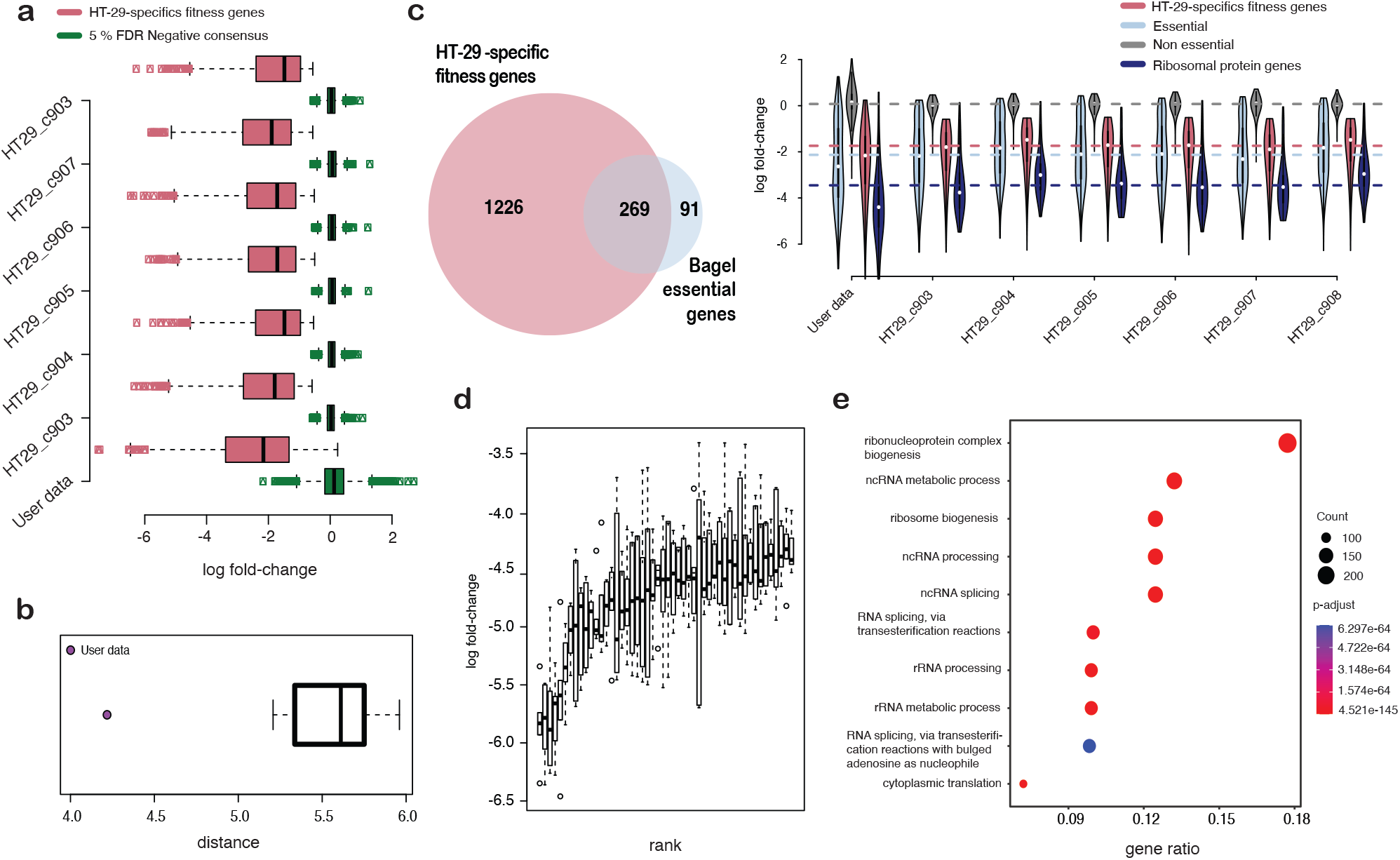
(a) Depletion log fold-change (LFCs) distributions of HT-29 specific positive and negative essential genes across individual reference HT-29 screens and example user-provided data. (b) Distribution of distances between HT-29 specific positive and negative essential genes, quantified through Cohen’s d, across the reference HT-29 screens (the boxplot) or for the example user-provided data (which in this case does not meet expectations). (c) On the left, comparing the HT-29-specific essential genes and a widely used set of prior known essential genes highlights a statistically significant overlap (two-sided Fisher’s exact test p-value = 7.1*x*10^−221^); on the right, distribution of LFCs for different gene sets along with the HT-29-specific fitness genes across the reference HT-29 screens, as well as an example user-provided data. (d) Depletion LFCs of the top 50 HT-29-specific fitness genes consistently depleted in all experiments, across HT-29 reference screens. (e) Top 10 Gene Ontology categories (Biological ProcessP) significantly enriched (Benjamini-Hochberg corrected p-value < 0.05) in the HT-29-specific essential genes.

### Usage Notes

The HT-29 reference dataset can be manually downloaded at https://score.depmap.sanger.ac.uk/downloads or http://iorio-apps.fht.org/, or on FigShare^17^. Alternatively, the function `HT29R.download_ref_dataset` of the *HT29benchmark* package can be used to download the reference dataset within an R session. A vignette with instructions on how to perform a quality assessment of a newly performed screen of the HT-29 cell line employing Sanger library^16^ and settings described in Behan *et al*.^7^ is provided with the package, together with a wrap-function that performs all the assessment steps and produces a final report in PDF format, as well as reproducing all the figures we presented here.

Users have a non-exclusive, non-transferable right to use data files for internal proprietary research and educational purposes, including target, biomarker and drug discovery. Excluded from this licence are the use of the data (in whole or any significant part) for resale either alone or in combination with additional data/product offerings, or for provision of commercial services. Both package and reference data are experimental and academic in nature and are not licensed or certified by any regulatory body. Furthermore, data access is provided on an “as is” basis and excludes all warranties of any kind (express or implied).

### Code availability

The R code used for generating this dataset, for its QC assessment, as well as to evaluate the quality of a user-provided screen and to reproduce all the figures presented here is available at https://github.com/francescojm/HT29benchmark.

## Acknowledgements

IM was supported by fellowships from the INCIPIT PhD program co-funded by Horizon 2020-COFUND Marie Sklodowska-Curie Actions, and an EMBO short-term fellowship (8536). This work has been partially funded by OpenTargets (OTAR-015 and OTAR2-055), and by Wellcome Trust Grant 206194. For the purpose of open access, the authors have applied a CC BY public copyright licence to any Author Accepted Manuscript version arising from this submission.

## Author contributions statement

RMI, FI, MJG conceived the study; RMI, IM, CP designed, and performed the benchmark analyses; RMI assembled the Jupyter notebook; RMI, AS, IM, CP, FI wrote and revised the manuscript; FB performed experiments underlying the HT-29 reference dataset under the supervision of MJG; RMI, IM, AS, interpreted results and assembled figures; MG and MJG contributed to study supervision; AS and FI supervised the study. All authors read and revised the manuscript.

