## Supplementary Information for "Benchmark data and software for assessing genome-wide CRISPR-Cas9 screening pipelines"

### Supplementary Figure 1

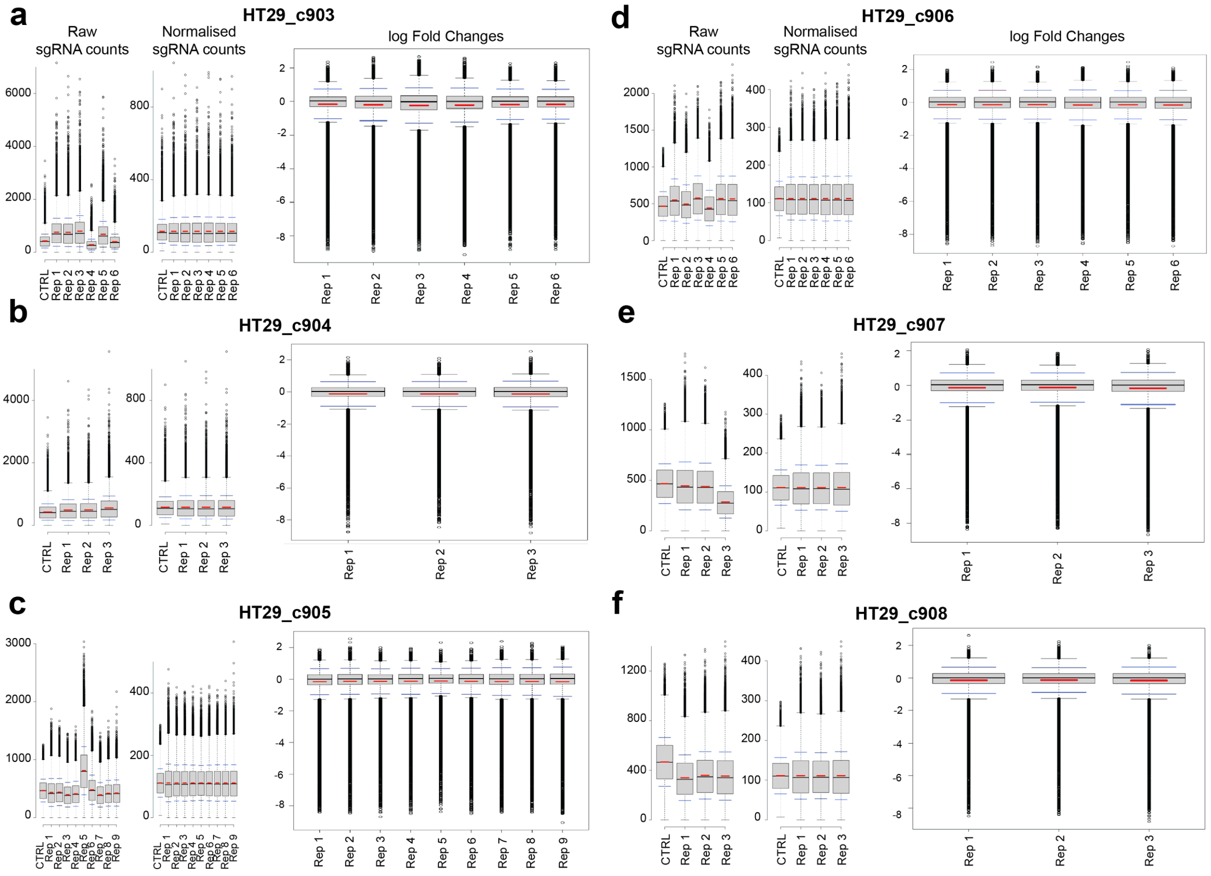


**Supplementary Figure 1: Normalised sgRNA counts and depletion log fold-changes across HT-29 screens.** Distributions of sgRNA counts in the plasmid library (CTRL) and the each HT-29 screen replicates 14 days post-selection before (leftmost plot) and after (centre plot) normalisation by total numbers of reads, and sgRNA depletion log fold-changes (one screen per each panel).

### Supplementary Figure 2

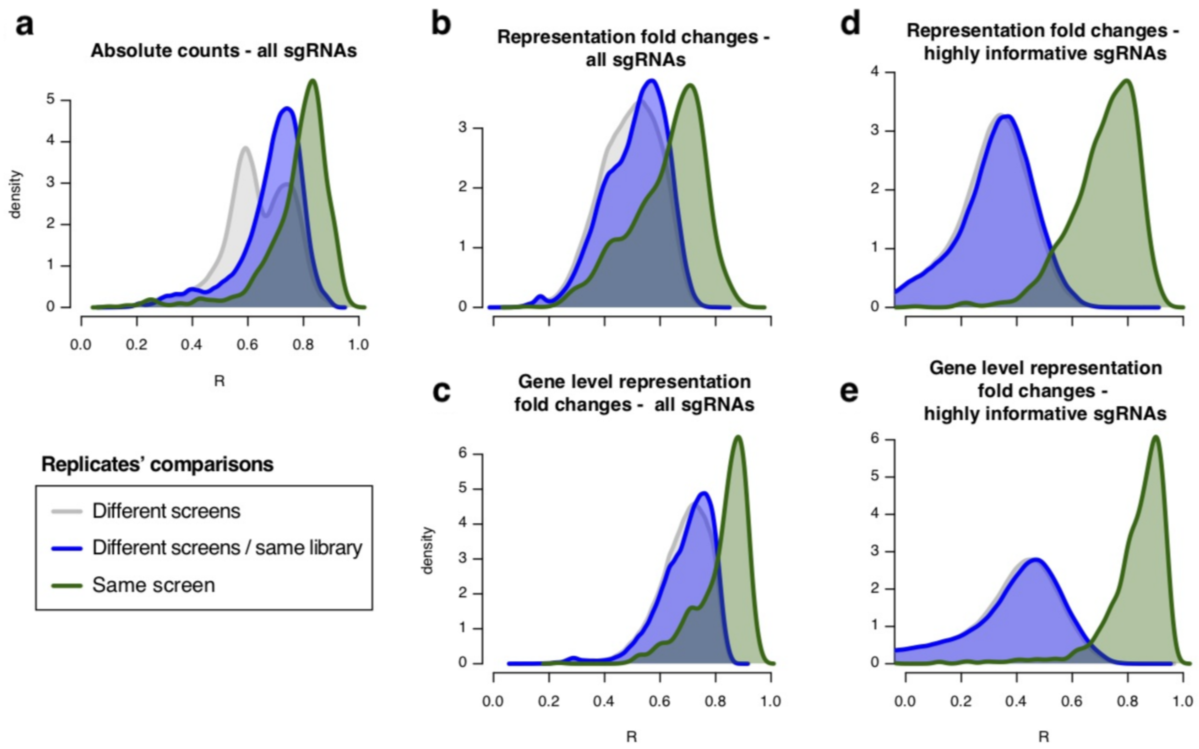


**Supplementary Figure 2: Distributions of Pearson's correlations between individual screen replicates from Project Score.** Computed (**a**) between profiles of sgRNA absolute counts, (**b**) log fold-changes (LFCs) of individual sgRNA aboundance 14 days post library transduction versus control plasmid, (**c**) as for B but with LFCs averaged on a targeted gene basis, (**d**) as for B but considering only highly informative/reproducible sgRNAs (defined in Behan et al, Nature 2019); (**e**) as for **c** but considering only gene highly informative/reproducible sgRNAs.

In ABC correlations between replicates of the same screens are not sufficiently distinguishable from correlations computed between individual replicates of different screens, thus Pearson's correlation cannot be used as a criteria to assess screens' reproducibility. In contrast, in DE correlations between replicates of the same screens are significantly larger than those between individual replicates of different screens, allowing the definition of a reproducibility threshold and the use of Pearson's correlation between replicates as a means to assess screens' reproducibility.

### Supplementary Figure 3

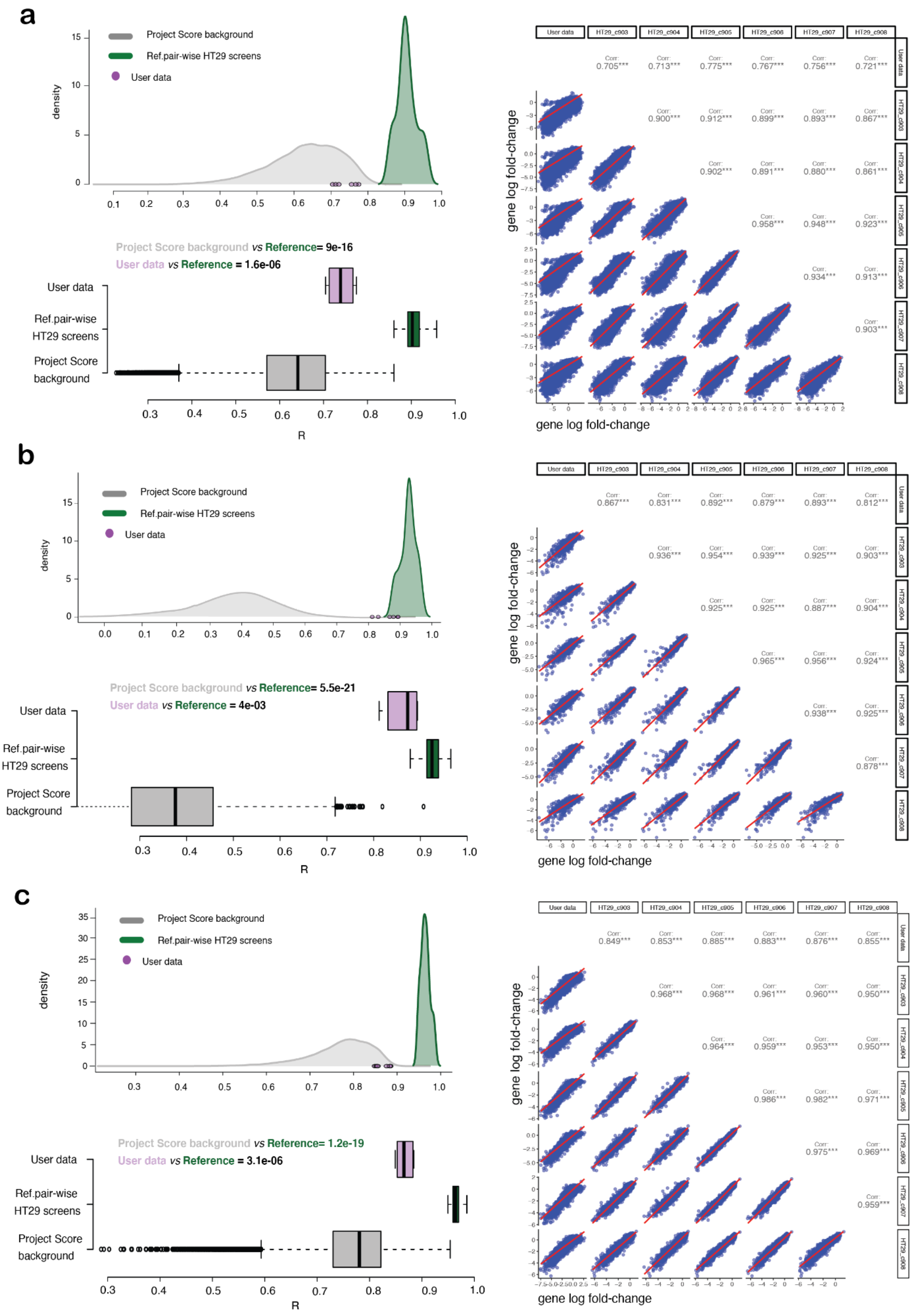

**Supplementary Figure 3: Distributions of Pearson's correlations between HT-29 screens and user-provided data.** Pearson's correlation scores between genome-wide profiles of depletion log fold-changes (LFCs) computed at the sgRNA (**a**) and at the gene level (**b**), or using only a subset of reproducible and highly informative sgRNAs (n = 838, **c**), between pairs of HT-29 screens (in green) and between the HT-29 reference screens and an example user-provided screen (in pink). The distributions in grey are computed between each possible pair of screen replicates in Project Score to estimate expectation. Two-sided t-test comparing expected Project Score correlation scores versus those computed between each pair of screens in the HT-29 reference dataset, as well as those computed between the example data screens versus those computed in the HT-29 reference dataset are also reported. The reference dataset scores are largely significantly different from expectation, the user data scores are still largely different from expectation but not as much as the reference data. The rightmost plots in each panel show Scatter-plot correlation matrices with pairwise Pearson's correlation scores computed within HT-29 reference screens and between user data versus HT-29 reference screens.

### Supplementary Figure 4

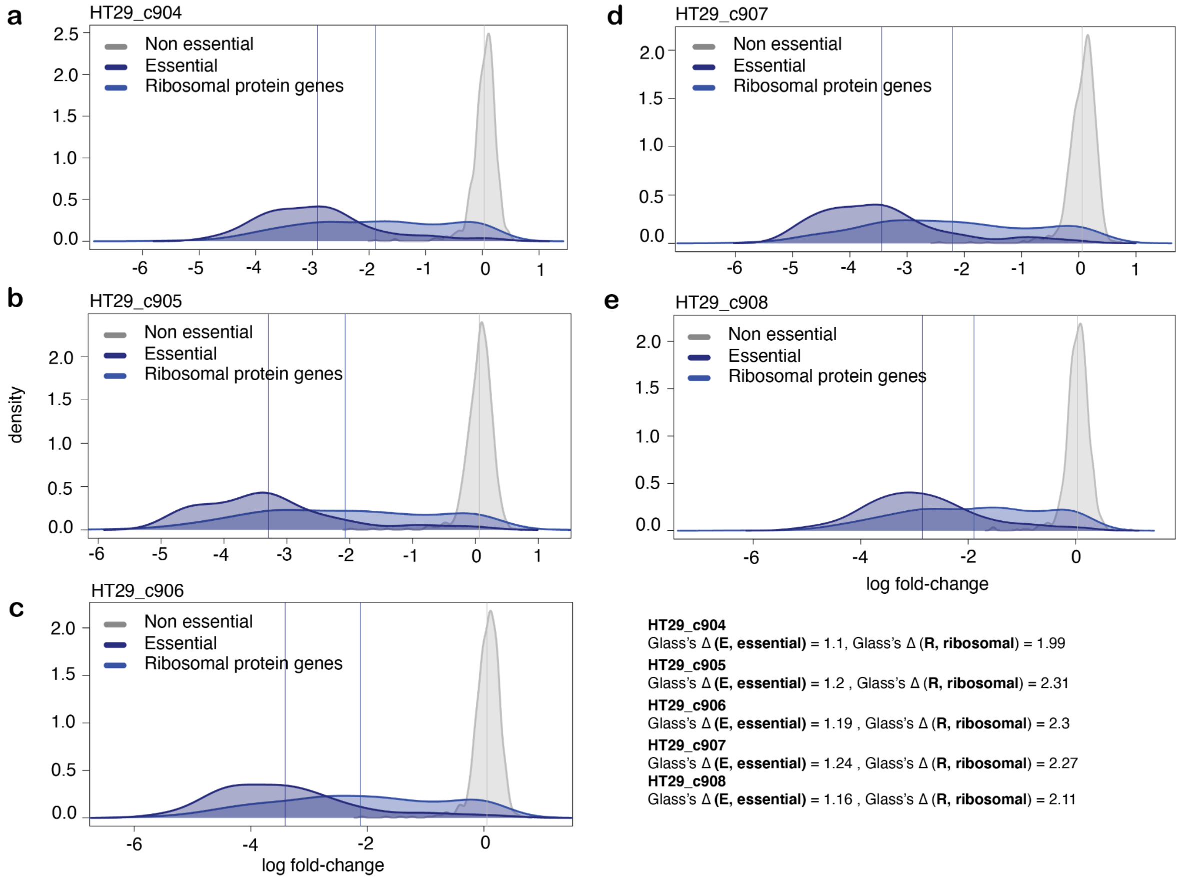


**Supplementary Figure 4: Reference HT-29 data screens quality in terms of phenotype Intensity.** Distributions of gene depletion log fold-changes (LFCs) for the screens in the HT-29 reference dataset (on screen for each panel). Glass's Δ (GD) scores for reference essential genes (*E*) and ribosomal protein genes (*R*) are also reported for each screen. Vertical lines indicate mean LFCs for each gene set as indicated by the different colours.

### Supplementary Figure 5

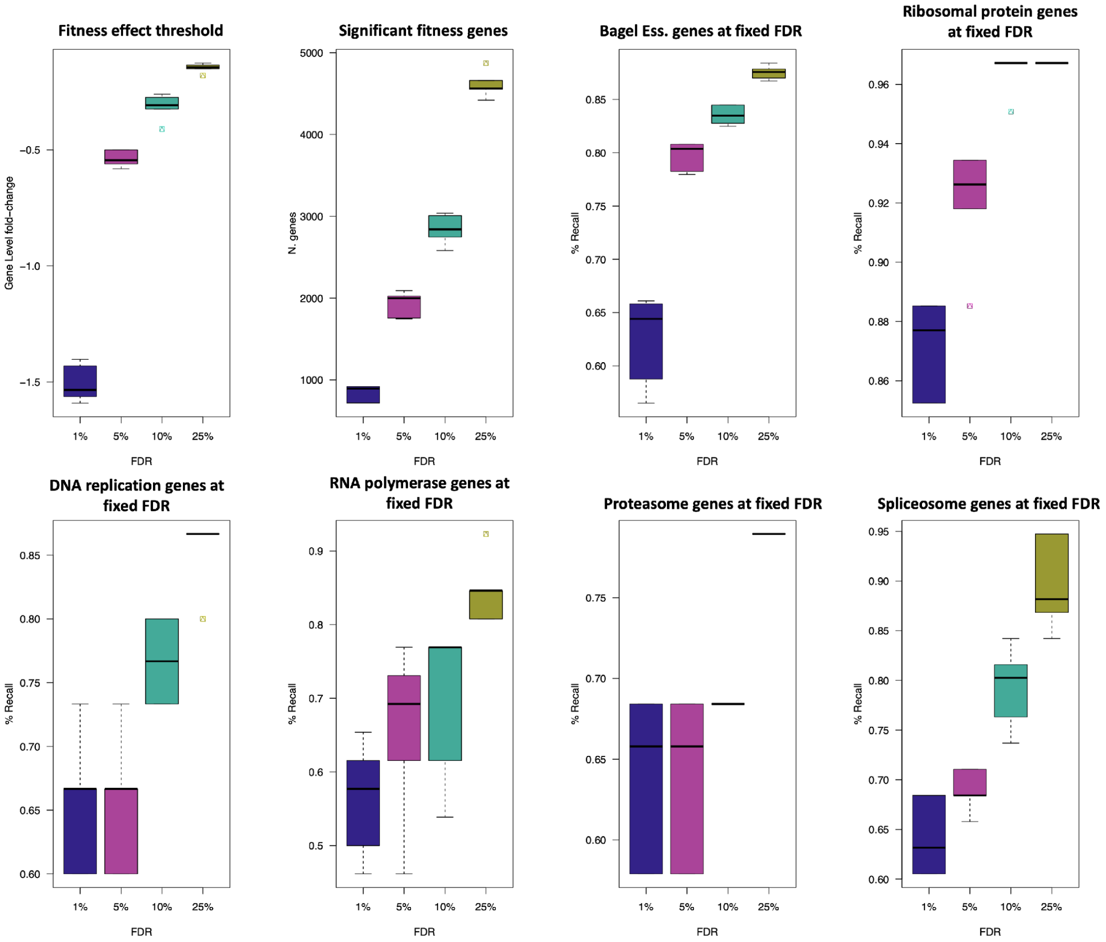


**Supplementary Figure 5.** Expected fitness effect significance threshold, number of fitness genes, and Recall of different sets of prior known essential genes at different levels of FDR computed using Bagel essential/ non-essential genes.

### Supplementary Table 1

| Screened model: HT-29 Colorectal carcinoma cell line, tissue: Large intestine, COSMIC ID: 905939, Cell Model Passports id: SIDM00136 | | | | | | | | |
|---|---|---|---|---|---|---|---|---|
| **Replicate identifier** | **Screen identifier** | **Sanger library** | **Cas9 Activity %** | **Transduction Efficiency (TE) %** | **Avg TE %** | **sgRNA count file name** | **Puromycin Selection Efficiency (PSE) %** | **Avg PSE** |
| HT29_c903R1 | HT29_c903 | Sanger V1.0 | 94.8 | 31.2 | 32.33 | HT29_c903R1.read_count.tsv.gz | 83.9 | 83.53 |
| HT29_c903R2 | | Sanger V1.0 | 94.8 | 36.6 | | HT29_c903R2.read_count.tsv.gz | 79.4 | |
| HT29_c903R3 | | Sanger V1.0 | 94.8 | 35.8 | | HT29_c903R3.read_count.tsv.gz | 84.8 | |
| HT29_c903R4 | | Sanger V1.0 | 94.8 | 30 | | HT29_c903R4.read_count.tsv.gz | 84.4 | |
| HT29_c903R5 | | Sanger V1.0 | 94.8 | 29.1 | | HT29_c903R5.read_count.tsv.gz | 84.4 | |
| HT29_c903R6 | | Sanger V1.0 | 94.8 | 31.3 | | HT29_c903R6.read_count.tsv.gz | 84.3 | |
| HT29_c904R1 | HT29_c904 | Sanger V1.0 | 94.8 | 29.2 | 27.57 | HT29_c904R1.read_count.tsv.gz | 90.1 | 89.97 |
| HT29_c904R2 | | Sanger V1.0 | 94.8 | 27 | | HT29_c904R2.read_count.tsv.gz | 89.4 | |
| HT29_c904R3 | | Sanger V1.0 | 94.8 | 26.5 | | HT29_c904R3.read_count.tsv.gz | 90.4 | |
| HT29_c905R1 | HT29_c905 | Sanger V1.1 | 94.8 | 26.8 | 33.42 | HT29_c905R1.read_count.tsv.gz | 82.4 | 80.81 |
| HT29_c905R2 | | Sanger V1.1 | 94.8 | 25.9 | | HT29_c905R2.read_count.tsv.gz | 80.2 | |
| HT29_c905R3 | | Sanger V1.1 | 94.8 | 26 | | HT29_c905R3.read_count.tsv.gz | 81.2 | |
| HT29_c905R4 | | Sanger V1.1 | 94.8 | 39.9 | | HT29_c905R4.read_count.tsv.gz | 74.5 | |
| HT29_c905R5 | | Sanger V1.1 | 94.8 | 38.1 | | HT29_c905R5.read_count.tsv.gz | 77.4 | |
| HT29_c905R6 | | Sanger V1.1 | 94.8 | 39.1 | | HT29_c905R6.read_count.tsv.gz | 76 | |
| HT29_c905R7 | | Sanger V1.1 | 94.8 | 37 | | HT29_c905R7.read_count.tsv.gz | 85.8 | |
| HT29_c905R8 | | Sanger V1.1 | 94.8 | 35.1 | | HT29_c905R8.read_count.tsv.gz | 86.4 | |
| HT29_c905R9 | | Sanger V1.1 | 94.8 | 32.9 | | HT29_c905R9.read_count.tsv.gz | 83.4 | |
| HT29_c906R1 | HT29_c906 | Sanger V1.1 | 94.8 | 30.1 | 35.65 | HT29_c906R1.read_count.tsv.gz | 89 | 88.40 |
| HT29_c906R2 | | Sanger V1.1 | 94.8 | 30.2 | | HT29_c906R2.read_count.tsv.gz | 88.3 | |
| HT29_c906R3 | | Sanger V1.1 | 94.8 | 29.4 | | HT29_c906R3.read_count.tsv.gz | 86.2 | |
| HT29_c906R7 | | Sanger V1.1 | 94.8 | 43.7 | | HT29_c906R7.read_count.tsv.gz | 89.8 | |
| HT29_c906R8 | | Sanger V1.1 | 94.8 | 40.8 | | HT29_c906R8.read_count.tsv.gz | 87.2 | |
| HT29_c906R9 | | Sanger V1.1 | 94.8 | 39.7 | | HT29_c906R9.read_count.tsv.gz | 89.9 | |
| HT29_c907R7 | HT29_c907 | Sanger V1.1 | 94.8 | 33.6 | 32.40 | HT29_c907R7.read_count.tsv.gz | 90 | 89.07 |
| HT29_c907R8 | | Sanger V1.1 | 94.8 | 30 | | HT29_c907R8.read_count.tsv.gz | 88.9 | |

| | | | | | | | | |
|---|---|---|---|---|---|---|---|---|
| HT29_c907R9 | | Sanger V1.1 | 94.8 | 33.6 | | HT29_c907R9.read_count.tsv.gz | 88.3 | |
| HT29_c908R4 | HT29_c908 | Sanger V1.1 | 94.8 | 32 | 32.00 | HT29_c908R4.read_count.tsv.gz | 74 | 79.33 |
| HT29_c908R5 | | Sanger V1.1 | 94.8 | 32 | | HT29_c908R5.read_count.tsv.gz | 73 | |
| HT29_c908R6 | | Sanger V1.1 | 94.8 | 32 | | HT29_c908R6.read_count.tsv.gz | 91 | |

**Supplementary Table 1.** Reference dataset manifest and experimental settings.

### Supplementary Table 2

| Fitness genes | 1% FDR | 5% FDR | 10% FDR | 25% FDR |
|---|---|---|---|---|
| Fitness effect threshold | -1.54 [-1.577, -1.480] | -0.55 [-0.571, -0.517] | -0.32 [-0.367, -0.284] | -0.146 [-0.164, -0.138] |
| Significant fitness genes | 875 [772.5, 915.0] | 1989 [1751.5, 2015] | 2756 [2664, 2965.5] | 4555 [4485.5, 4616] |
| Recall of E genes | 0.634 [0.600, 0.655] | 0.799 [0.781, 0.807] | 0.830 [0.826, 0.841] | 0.872 [0.868, 0.878] |
| Recall of Ribosomal protein genes | 0.868 [0.852, 0.885] | 0.918 [0.918, 0.934] | 0.967 [0.959, 0.967] | 0.967 [0.967, 0.967] |
| Recall of DNA replication genes | 0.666 [0.600, 0.666] | 0.666 [0.600, 0.666] | 0.733 [0.733, 0.800] | 0.866 [0.866, 0.866] |
| Recall of RNA polymerase genes | 0.577 [0.500, 0.596] | 0.653 [0.557, 0.730] | 0.769 [0.673, 0.769] | 0.846 [0.807, 0.846] |
| Recall of Proteasome genes | 0.631 [0.578, 0.684] | 0.631 [0.578, 0.684] | 0.684 [0.684, 0.684] | 0.789 [0.789, 0.789] |
| Recall of Spliceosome genes | 0.631 [0.605, 0.657] | 0.684 [0.671, 0.697] | 0.789 [0.750, 0.815] | 0.868 [0.855, 0.921] |

**Supplementary Table 2.** Reference table with median and interquartile range for expected fitness effect significance threshold, number of fitness genes, and recall of different sets of prior known essential genes, across different fixed levels of FDR (False Discovery Rate).
